# Personalized Pangenome References

**DOI:** 10.1101/2023.12.13.571553

**Authors:** Jouni Sirén, Parsa Eskandar, Matteo Tommaso Ungaro, Glenn Hickey, Jordan M. Eizenga, Adam M. Novak, Xian Chang, Pi-Chuan Chang, Mikhail Kolmogorov, Andrew Carroll, Jean Monlong, Benedict Paten

## Abstract

Pangenomes, by including genetic diversity, should reduce reference bias by better representing new samples compared to them. Yet when comparing a new sample to a pangenome, variants in the pangenome that are not part of the sample can be misleading, for example, causing false read mappings. These irrelevant variants are generally rarer in terms of allele frequency, and have previously been dealt with using allele frequency filters. However, this is a blunt heuristic that both fails to remove some irrelevant variants and removes many relevant variants. We propose a new approach, inspired by local ancestry inference methods, that imputes a personalized pangenome subgraph based on sampling local haplotypes according to *k*-mer counts in the reads. Our approach is tailored for the Giraffe short read aligner, as the indexes it needs for read mapping can be built quickly. We compare the accuracy of our approach to state-of-the-art methods using graphs from the Human Pangenome Reference Consortium. The resulting personalized pangenome pipelines provide faster pangenome read mapping than comparable pipelines that use a linear reference, reduce small variant genotyping errors by 4x relative to the Genome Analysis Toolkit (GATK) best-practice pipeline, and for the first time make short-read structural variant genotyping competitive with long-read discovery methods.

## 1 Introduction

Pangenome graphs [7] represent aligned haplotype sequences. Each node in the graph is labeled with a sequence, and each path represents the sequence obtained by concatenating the node labels. While any path in the graph is a potential haplotype, most paths are unlikely recombinations of true haplotypes.

One common application for pangenome graphs is as a reference for read mapping [11, 28, 31]. Graph-based aligners tend to be more accurate than aligners using a linear reference sequence when mapping reads to regions where the sequenced genome differs from the reference sequence. If the graph contains another haplotype close enough to the sequenced genome in that region, the aligner can usually find the correct mapping. On the other hand, sequence variation that is present in the graph but not in the sequenced genome can make the aligner less accurate. Such variation can imitate other regions, making incorrect mappings more likely.

Because having the right variants in the graph has a greater effect on mapping accuracy than having variants not present in the sequenced genome, the usual approach is building a graph based on common variants only. The vg short read aligner [11] got best results with the *1000 Genomes Project* (1000GP) [33] graph when variants below 1% threshold were left out. Pritt et al. [26] developed a model for selecting variants based not only on their frequency but also on their effect on the repetitiveness of the reference.

The Giraffe short read aligner [31] uses another approach for avoiding rare variants. As a haplotype-aware aligner, Giraffe can generate synthetic haplotypes by sampling local haplotypes proportionally with a greedy algorithm. It got best results with the 1000GP graph by generating 64 haplotypes. However, frequency filtering was still used with the *Human Pangenome Reference Consortium* (HPRC) [20] subgraph intended for Giraffe. As the initial graphs contain only 90 haplotypes, the threshold was set to 10%.

Vaddadi et al. [34] have proposed a VCF-based pipeline for building a personalized diploid reference genome. They genotype variants in a reference VCF using subsampled reads and impute the haplotypes based on the reference panel. The resulting personalized VCF file can be used for building a graph for any graph-based read mapping and variant calling pipeline.

In this paper, we propose a more direct approach with less overhead. We create a *personalized pangenome reference* by sampling haplotypes that are similar to the sequenced genome according to *k*-mer counts in the reads. We work directly with assembled haplotypes, and the sampled graph we build is a subgraph of the original graph. Therefore any alignments in the sampled graph are also valid in the original graph.

Our approach is tailored for Giraffe, as the indexes it needs for read mapping can be built quickly. We assume a graph with a linear high-level structure, such as graphs built using the Minigraph– Cactus pipeline [13]. We further assume that read coverage is high enough (at least 20x) that we can reliably classify *k*-mers into absent, heterozygous, and homozygous according to *k*-mer counts.

When used with HPRC graphs, our sampling approach increases the overall running time of the pipeline by less than 15 minutes. We show that the sampled graph is a better mapping target than the universal frequency-filtered graph. We see a small improvement in the accuracy of calling small variants with DeepVariant [24] and a large increase in the accuracy of genotyping structural variants (SVs) with vg [12] and PanGenie [6].

## 2 Methods

See Figure 1 for an overview of the methods; the following describes the method in detail.

**Figure 1:**
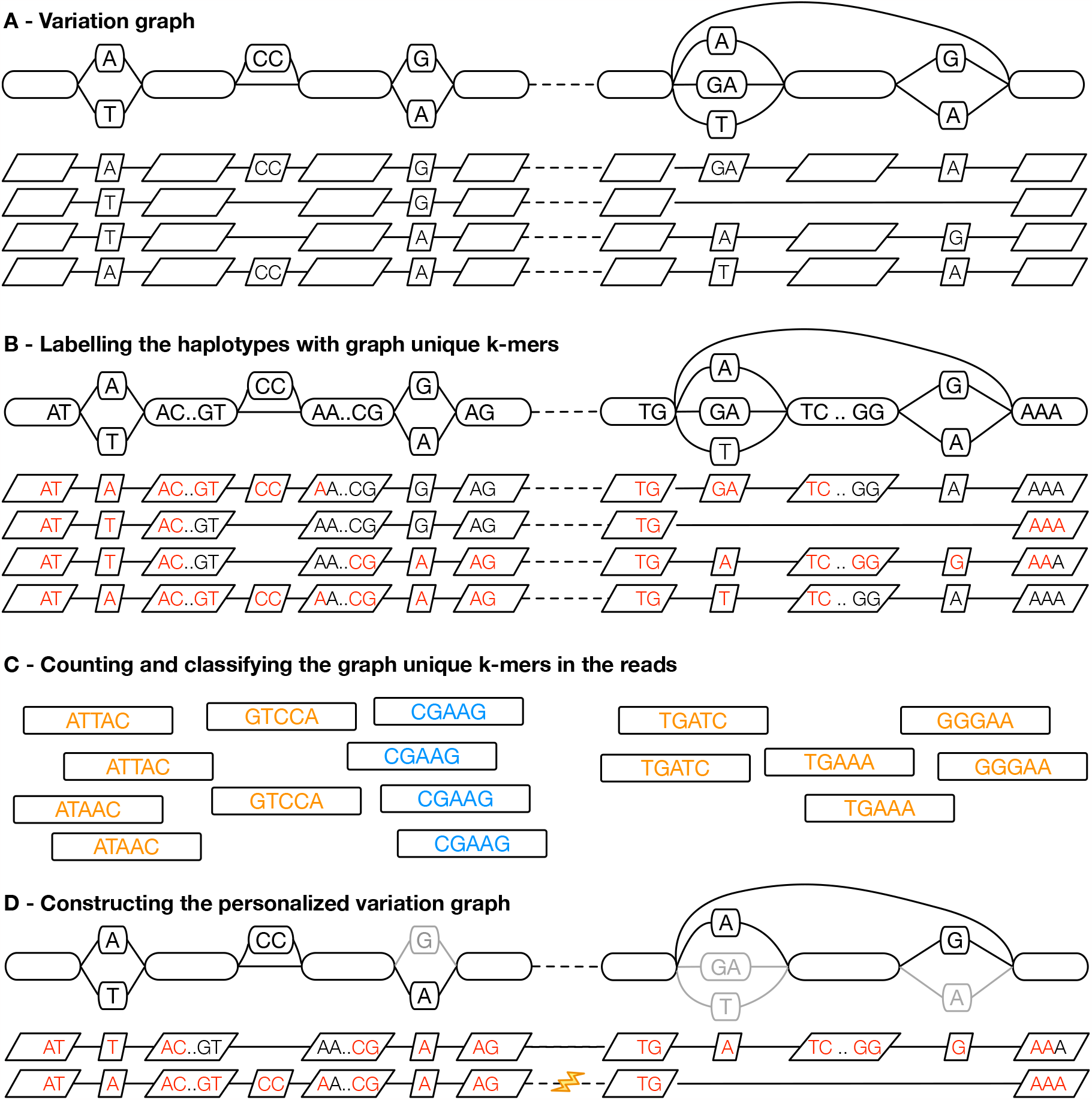
Illustrating haplotype sampling at adjacent blocks in the pangenome. (A) A variation graph representing adjacent locations in the pangenome, composed of a bidirected sequence graph (top) and a set of embedded haplotypes (below); the dotted lines represent the boundary between the two blocks. (B) *k*-mers that occur once within the graph, termed graph-unique k-mers, are identified in the haplotypes; here *k* = 5 and graph-unique *k*-mers are colored red. The presence and absence of these graph-unique *k*-mers identifies each haplotype. (C) The graph-unique *k*-mers are counted in the reads, and based upon counts classified as present, likely heterozygous (shown in orange), present, likely homozygous (shown in blue), or absent (all red kmers in (B) not identified in the reads). (D) Using the identified graph-unique *k*-mer classifications, a subset of haplotypes are selected at each location, defining a personalized pangenome reference subgraph of the larger graph. Where needed, recombinations are introduced (see lightning bolt) to create contiguous haplotypes.

### 2.1 Bidirected sequence graphs

Human pangenome graphs are usually based on the *bidirected sequence graph* model. Nodes have identifiers, and they contain a sequence. Edges are undirected and connect two node sides. A forward traversal of a node enters from the left, reads the sequence, and exits from the right. A reverse traversal enters from the right, reads the reverse complement of the sequence, and exits from the left.

Bidirected sequence graphs can be stored in the text-based *Graphical Fragment Assembly* (GFA) format. The *GBZ file format* [29] is a space-efficient binary format for pangenome graphs. It is based on the GBWT index [30] that stores haplotype paths as node sequences. GBZ is compatible with a subset of GFA, and the two can be converted efficiently to each other.

The structure of a bidirected sequence graph can be described hierarchically by its *snarl decomposition* [23]. A snarl is a generalization of a bubble [38], and denotes a site of genomic variation. It is a subgraph separated by two node sides from the rest of the graph. Each snarl must be minimal in the sense that neither of the sides defining it forms a snarl with a node side located within the subgraph. A graph can be decomposed into a set of chains, each of which is a sequence of nodes and snarls. A snarl may either be primitive, or it may be further decomposed into a set of chains.

### 2.2 Preprocessing haplotypes

We assume that the graph is in the GBZ format. While the format is space-efficient and supports efficient queries, it is not suitable for selecting haplotypes based on sequence similarity. We therefore need to preprocess the graph and store the haplotypes in a more appropriate format.

Our preprocessing approach is similar to that used in PanGenie [6] (Figure 1A). We assume that each weakly connected component in the graph corresponds to a single top-level chain in the snarl decomposition of the graph. While PanGenie combines bubbles that are less than *k* bp apart (default *k* = 31), we combine adjacent snarls in the top-level chains into approximately *b* bp *blocks* (default *b* = 10000), using a minimum distance index [3] for determining the length of a block.

Top-level chains generally correspond to chromosomes or other meaningful contigs. Because we partition each top-level chain into a sequence of blocks, we can ensure that sampled haplotype paths corresponding to that contig visit the same blocks in the same order. If we sample the haplotypes independently in each block, we can therefore assume that any recombination of them is a plausible haplotype. Despite this linear high-level structure, the graph may contain reversals that let the same real haplotype visit the same blocks multiple times. To avoid sampling unbalanced recombinations, we consider each minimal end-to-end visit to a block a separate haplotype in that block. We find such visits efficiently by listing the paths visiting the border nodes of a block using the r-index [10] and matching the node visits at both ends of the block.

We describe the haplotypes in each block in terms of *k*-mers that are specific to the block (Figure 1B). While PanGenie uses *k*-mers (default *k* = 31) with at most a single hit in each haplotype (but possibly multiple hits within the bubble), we use minimizers (default *k* = 29, *w* = 11) with a single hit in the graph (but possibly multiple hits in a haplotype). We also avoid uninformative *k*-mers that are present in each haplotype. For each block, we then build a *k-mer presence matrix* : a binary matrix that marks the presence/absence of each selected graph-unique *k*-mer in each haplotype.

### 2.3 Sampling haplotypes

The haplotype sampling workflow starts with counting *k*-mers in the reads (Figure 1C). Any external tool that supports the *K-mer File Format* (KFF) [5] can be used. The sampling algorithm ignores *k*-mers with a single occurrence, and it combines the counts of a *k*-mer and its reverse complement. It also ignores all *k*-mers not used in the *k*-mer presence matrices.

If *k*-mer coverage is not provided, we estimate it from the counts. If the most common count is above the median, we use it as the estimate. Otherwise, if there is a good enough secondary peak above the median at approximately 2x the primary peak, we use it. If both attempts fail, the user must provide an estimate. Given the *k*-mer coverage and *k*-mer counts, we classify each *k*-mer in the matrices as absent, heterozygous, homozygous, or frequent. We then ignore all frequent *k*-mers. While this classification is inherently noisy, on aggregate, it is useful for our purposes.

We sample the haplotypes independently in each block (Figure 1D). This is in contrast to PanGenie, which attempts to infer phased haplotypes. Sampling is based on a greedy algorithm that selects the highest-scoring haplotype and then adjusts the scores for the remaining haplotypes. The score of a haplotype is the sum over *k*-mer scores. Initially each homozygous *k*-mer yields +1 or −1, depending on if it is present or absent in the haplotype. Absent *k*-mers get −0.8 or +0.8, respectively, while heterozygous *k*-mers do not contribute to the initial scores. (See Supplement 2 for a discussion on how the sampling parameters were selected.)

The first haplotype we select is an approximation of the consensus sequence. With the subsequent haplotypes, we aim to cover all homozygous *k*-mers while selecting haplotypes both with and without heterozygous *k*-mers. We discount the scores of homozygous *k*-mers that were present in the selected haplotype by a multiplicative factor 0.9. With heterozygous *k*-mers, we adjust the scores by an additive term 0.05 in the other direction, making it more likely that the next haplotype is the opposite with respect to the presence of this *k*-mer.

We can use the selected *n* haplotypes directly (typically *n* = 4 to 32), or we may use them as candidates in optional *diploid sampling*. In diploid sampling, we consider each pair of candidates separately and select the highest-scoring pair. For each *k*-mer, we determine the number of copies *x* (0, 1, or 2) we expect based on *k*-mer counts and the number of copies *y* that are present in the candidates. The score for that *k*-mer is then 1 − |*x* − *y*|, and the score for the pair is the sum of *k*-mer scores.

After sampling local haplotypes in each block of a top-level chain, we combine them to form full-length haplotypes. If the same haplotype was selected in adjacent blocks, we connect them together. Otherwise, the haplotypes we form are arbitrary recombinations of the local haplotypes we sampled. We insert the haplotypes into an empty GBWT index, along with any reference paths for that chain if we want to include them in the personalized pangenome reference.

The sampling process can be parallelized over the top-level chains. Because top-level chains correspond to weakly connected components in the graph, the resulting GBWT indexes are disjoint and can be merged efficiently. Once the final GBWT is made, we build the GBZ graph induced by the haplotypes in the index. Because this *sampled graph* is a subgraph of the original graph, any alignment in it is also valid in the original graph.

## 3 Results

See Supplement 1 for further details on the exact data used.

### 3.1 HPRC Pangenome Graphs

The experiments below feature pangenome graphs constructed from the HPRC draft pangenome dataset. They were all generated with Minigraph-Cactus [13] using the same 90 haplotypes (44 diploid samples, GRCh38 and T2T-CHM13), whose assemblies are described in detail in [20]. The graphs were referenced on GRCh38, meaning this genome was left acyclic and unclipped by Minigraph-Cactus. The needs of the present study in part drove the creation of a v1.1 release of these graphs [14], created with a newer version of the tools, which is the primary version used here. The majority of changes to the graph construction methodologies used are detailed in [13]. One change that was not previously described but is relevant for this work is the implementation of a filter that guarantees that each connected component has a single top-level chain. This logic, implemented in vg clip and activated in Minigraph-Cactus as of v2.6.3, repeatedly removes all nodes which have one side of degree 0 (no edges attached) and that are not contained in the reference path, until none are left. Any edges on the left side of the first node of a reference path or the right side of the last node of the reference path are likewise removed. The resulting graph has exactly two “tips” (node sides with degree 0) per connected component, corresponding to the two ends of the reference chromosome. The snarl decomposition uses these two tips to root the snarl tree, guaranteeing a single top-level chain that can be used to guide the haplotype sampling algorithm described above. The number of nodes, tips, edges, and edges removed by this procedure, as well as its impact on the number of top-level chains, is shown in Supplementary Table 3. Allele frequency filtered versions of each graph were obtained from Minigraph-Cactus using a filter threshold of 9 (ie 10% of haplotypes). Sampling was only run on the unfiltered (default) graphs. Control results are presented for both the v1.1 graphs and the v1.0 graphs that were released alongside the HPRC data [20]. Since the v1.0 graphs have many top level chains per component, they cannot be used with the haplotype sampling approach described in this paper.

### 3.2 Mapping reads

The following benchmarks were done on an AWS i4i.16xlarge instance with 32 physical / 64 logical CPU cores and 512 GiB memory. We used a development version of vg that was effectively the same as version 1.52.0. All tools were allowed to use 32 threads.

We aligned 30x NovaSeq reads for the *Genome in a Bottle* (GIAB) HG002 sample to various references using Giraffe [31] and measured the running time. Note, the constructed pangenomes do not include any GIAB sample or close relative. We also used BWA-MEM [18] with a linear reference (GRCh38) as a baseline. For haplotype sampling, we first had to create a sampled graph for HG002. We counted *k*-mers in the reads using KMC [16]. Then we took the v1.1 default graph, containing all haplotypes, and ran haplotype sampling with 4, 8, 16, and 32 haplotypes as well as diploid sampling from 32 candidates. Before mapping the reads, we also had to build a distance index and a minimizer index for the sampled graph; these steps are now integrated into Giraffe to be run automatically.

The results can be seen in Figure 2. Giraffe is faster with all graphs than BWA-MEM is with a linear reference. Mapping the reads to the v1.1 filtered graph took less time than mapping them to the v1.0 filtered graph; anecdotally we observe that faster-to-compute alignments are often of higher quality. Giraffe was even faster with the v1.1 diploid graph, but the other steps (*k*-mer counting 432 s, haplotype sampling 341 s, index construction 997 s) increased the overall running time to the same level as with the v1.0 filtered graph. For sampled graphs without diploid sampling, both mapping time and the overall time increased with the number of haplotypes.

**Figure 2:**
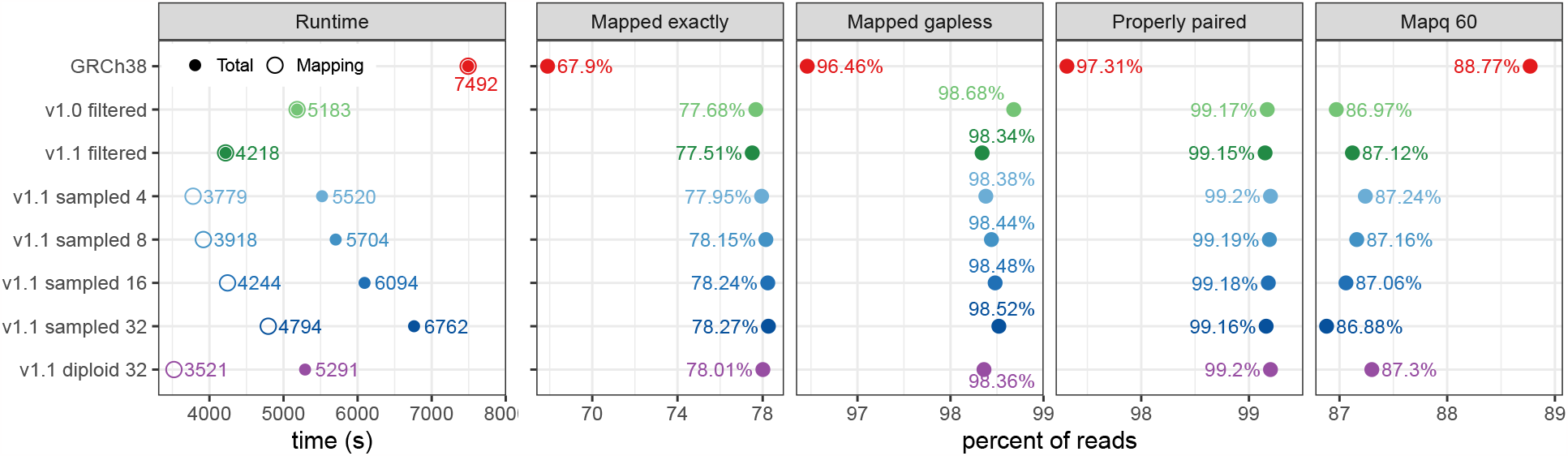
Mapping 30x NovaSeq reads for HG002 to GRCh38 (with BWA-MEM) and to HPRC graphs (with Giraffe). The graphs (y-axis) are Minigraph–Cactus graphs built using GRCh38 as the reference. For the sampled graphs, we tested sampling 4, 8, 16, and 32 haplotypes. For the v1.1 diploid graph, 32 candidate haplotypes were used for diploid sampling. We show the overall running time and the time spent for mapping only (left), and the fraction of reads with an exact, gapless, properly paired, and mapping quality 60 alignment.

Figure 2 also includes some statistics on the mapped reads. Giraffe is more likely to find an exact (i.e. all-matches) alignment with the v1.1 diploid graph than with the frequency-filtered graphs, and the reads are more likely to be properly paired. It is also more confident about the alignments to the v1.1 diploid graph, as evidenced by the higher proportion of reads with mapping quality (Mapq) 60. The variant calling and genotyping results in the following sections show this confidence is well-founded.

When we use sampled graphs without diploid sampling, we find more exact and gapless alignments than with the diploid graph, and the number of such alignments increases with number of haplotypes. On the other hand, we find fewer properly paired alignments, and a smaller fraction of the alignments get mapping quality 60. This indicates that while the additional haplotypes enable us to find better alignments, some of these alignments are likely wrong.

### 3.3 Calling small variants

We evaluated the impact of haplotype sampling on the accuracy of calling small variants, using the GIAB v4.2.1 GRCh38 benchmark set [35]. We mapped PCR-free NovaSeq 40x reads [1] with Giraffe [31] and called variants using DeepVariant [24], using the pipeline discussed in Liao et al. [20]. We compared the performance of different graphs, including the v1.1 filtered graph and v1.1 sampled graphs with 4, 8, 16, and 32 haplotypes, as well as the v1.1 diploid graph (Figure 3(A); Supplementary Table 5). The v1.1 diploid graph consistently outperformed the other graph configurations, reducing total errors by 9.1% across samples HG001 to HG005 relative to the v1.1 filtered graph. The performance of the v1.0 and v1.1 filtered graphs was similar (Supplementary Table 5). Sampling either 4 or 8 haplotypes improved performance relative to the filtered graphs, although we observe an inverse correlation between sampled haplotype number and performance.

**Figure 3:**
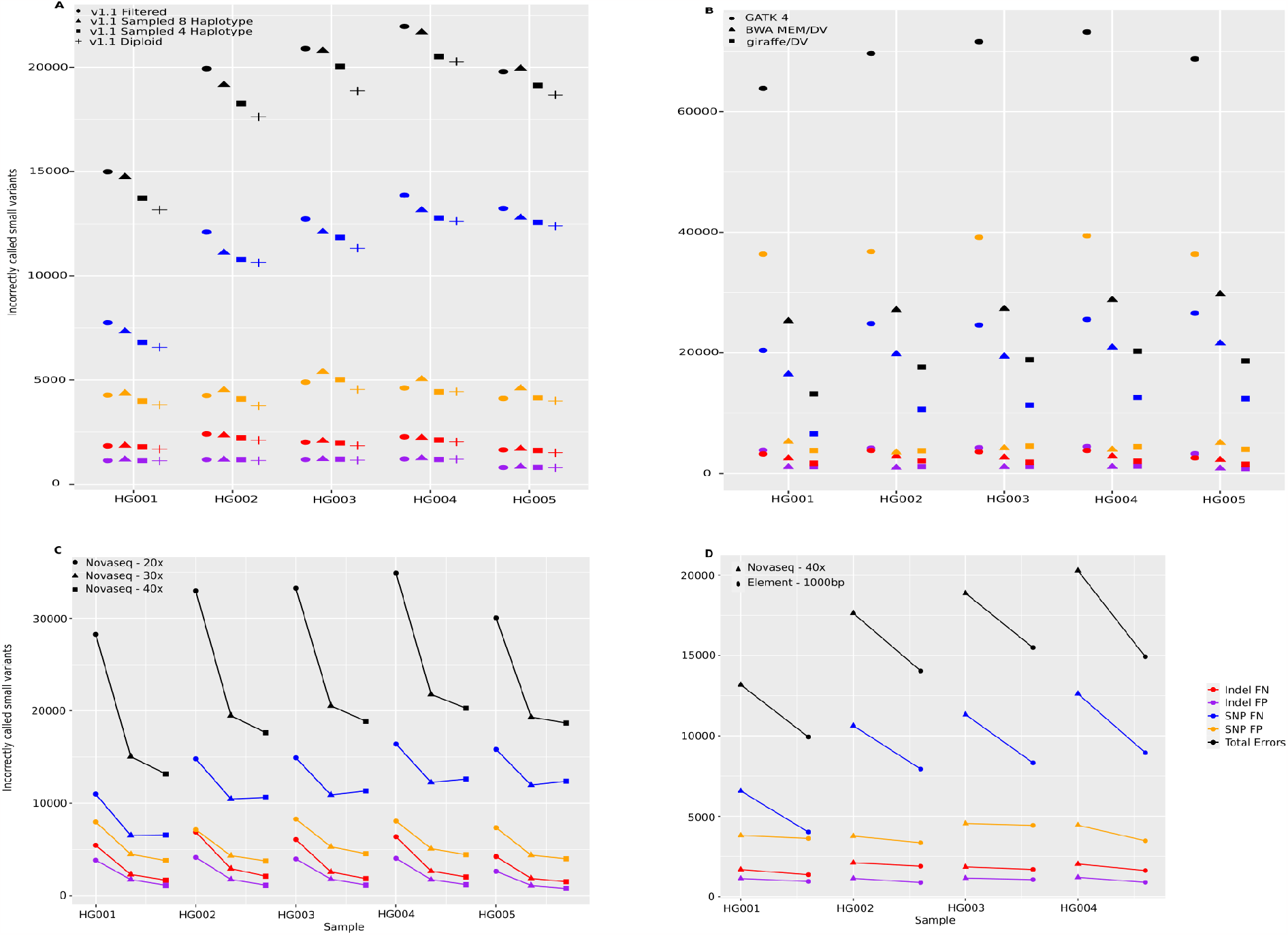
Small Variants evaluation across samples HG001 to HG005. (A) The number of false positive (FPs) and false negative (FNs) indels and SNPs across four different graphs, each using GRCh38 as the reference: v1.1 filtered, v1.1 sampled with 4 and 8 haplotypes, and v1.1 diploid, using the Giraffe/DeepVariant pipeline. (B) Comparing the Giraffe/DeepVariant using the v1.1 diploid graph to BWA MEM/DeepVariant and GATK best practice pipelines, both using the GRCh38 reference. (C) The performance of the Giraffe/DeepVariant pipeline using the v1.1 diploid graph with different coverage levels of NovaSeq reads (20x, 30x, and 40x). (D) Comparing the number of errors using either NovaSeq 40x data or Element 36x - 1000bp insert data; in both cases, using the Giraffe/DeepVariant pipeline with the v1.1 diploid graph. HG005 Element sequencing data was not available for comparison.

We compared performance to mapping to a linear reference (GRCh38) with BWA-MEM [18] and using DeepVariant to call variants, as well as to the GATK best-practice [25] pipeline (Figure 3(B)). Mapping the same NovaSeq 40x data to the v1.1 diploid graph with Giraffe reduced errors by an average 35% relative to BWA-MEM/GRCh38, and 74% (a near four-fold reduction in total errors) relative to the GATK best-practice.

We examined the effect of varying read coverage on the performance of the v1.1 diploid graph (Figure 3(C)). We observed large improvements in increasing coverage from 20 to 30x (avg. 39.7% reduction in errors), and more minor improvements going from 30 to 40x (avg. 7.8% reduction). It is worth stating that for all samples Giraffe/DV with 20x data made substantially fewer errors than GATK best-practice with 40x data.

To determine how a different read technology would affect the results, we repeated the analysis with sequencing data from Element Biosciences [2] (Figure 3(D), Supplementary Table 5). Relative to the comparable coverage Illumina dataset (Illumina NovaSeq 40x, Element 36x 1000 bp insert), we find a net reduction in errors of 22%, recapitulating the prior Google/Element results [2] showing this Element data to be of excellent quality.

### 3.4 Genotyping structural variants

We assessed the ability of pangenome SVs genotyping algorithms (PanGenie [6] and `vg call` [12]) using different pangenome graphs. We used the GIAB Tier1 v0.6 truth set comprising the set of confident structural variants for the HG002 sample. Haplotype sampling improves genotyping performance relative to both the v1.1 default and v1.1 filtered graphs (Figure 4(A); Supplementary Table 6); the combination of the v1.1 graphs and haplotype sampling also substantially improves upon the performance of 1.0 graphs described in Liao et al. [20] for the draft human pangenome. For example, using PanGenie, the F1 score of variant calls increased from 0.7780 using the v1.0 filtered graph to 0.8926 with the v1.1 diploid graph.

**Figure 4:**
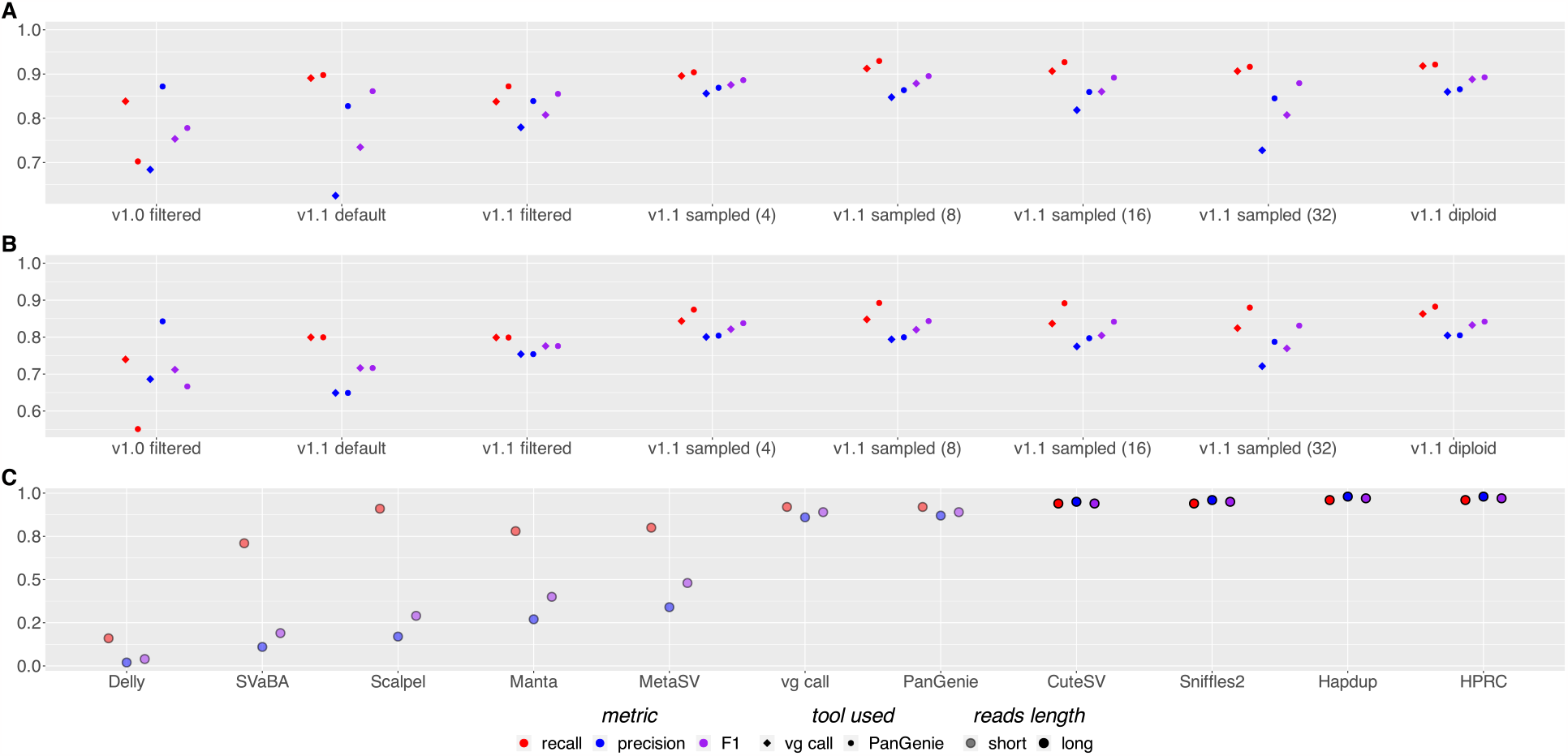
SVs benchmark evaluation. (A) Precision, recall, and F1 scores of both `vg call` and PanGenie for different pangenome reference graphs on the GIAB v0.6 Tier1 call set. Graphs were built using GRCh38 as the reference. (B) As with (A) but using a benchmark set of SVs created with DipCall from the T2T v0.9 HG002 genome assembly, comparing genome-wide but excluding centromeres. (C) Comparing the performance of PanGenie and `vg call` using the 1.1 diploid graph to other genotyping methods. Illumina short reads were used with Delly [27], SVaBA [37], Scalpel [8], Manta [4] and MetaSV [22] as well as with `vg call` [12] and PanGenie [6]. Also shown are long-read methods (CuteSV [15], Sniffles2 [32], Hapdup [17], and Human Pangenome Reference Consortium de novo assemblies [20]).

Regarding the number of haplotypes sampled, as with small variants, increasing beyond 8 hinders genotyping performance, with v1.1 diploid graph performing overall best. In general, PanGenie outperforms `vg call` across different graph builds and sampling strategies, although the difference between `vg call` and PanGenie for the best performing graph (v1.1 diploid graph) is small (F1 of 0.8926 for PanGenie vs. 0.888 for `vg call`). `vg call` also has an average 30.53% faster runtime (Supplementary Table 4 and Supplementary Figure 3).

We tested the performance of haplotype sampling using the Challenging Medically Relevant Genes (CMRG) benchmark (see Supplementary Tables 7 and 9) from GIAB [36]. We find similar trends to those observed in the whole genome, but performance drops slightly given the difficult nature of these regions. For example, the best-performing method in this benchmark, `vg call`, with the 1.1 diploid graph has an F1 score of 0.8683, whereas using the v0.6 GIAB benchmark its F1 score is 0.888.

The GIAB benchmark sets cover a limited subset of SVs in the genome. To get a more complete assessment of performance, we benchmarked our approach using a call set derived using DipCall [19] from a draft Telomere-to-Telomere (T2T) assembly of the HG002 sample (Figure 4(B); Supplementary Table 8). These assemblies are available on AWS at the following link https://s3-us-west-2.amazonaws.com/human-pangenomics/T2T/HG002/assemblies/hg002v0.9.fasta.gz, and more details on the process to obtain them and ongoing work towards a HG002 QV100 assembly set can be found at Telomere-to-telomere consortium HG002 “Q100” project. Excluding centromeric regions, which can not yet be reliably compared, we generate a call set covering 92.02% of the GRCh38 reference containing 25732 variants; in contrast the GIAB Tier1 v0.6 call set contains 9575 variants covering 86.29% of the reference. Across this larger call set, we find similar performance patterns to the GIAB v0.6 call set, with an F1 score of 0.8419 genome-wide, excluding centromeres, for PanGenie with the v1.1 diploid graph.

We finally compared our haplotype sampling results with other methods using the GIAB Tier1 v0.6 SVs benchmark set (Figure 4(C)). Relative to contemporary methods using short reads and the GRCh38 reference, our results with the v1.1 diploid graph are dramatically better (the F1 score for the best-performing short-read method, MetaSV is 0.29, whereas PanGenie achieves 0.89), and are now close to rivaling those using long reads.

## 4 Discussion

We have developed a *k*-mer-based method for sampling local haplotypes that are similar to a sequenced genome. When a subgraph based on the sampled haplotypes is used as a reference for read mapping, the results are more accurate than with the full graph or with a frequency-filtered graph containing only common variants. This translates into a small improvement in the accuracy of calling small variants and a large improvement in the accuracy of genotyping structural variants relative to existing pangenome approaches. While the presented results represent an advance over previous filter-based methods, several limitations exist.

Our *k*-mer presence matrices are not compressed. With the HPRC v1.1 graph in Section 3.2, we used 176 million *k*-mers for describing the haplotypes. This corresponds to 21 MiB per haplotype, plus overhead. As the number of haplotypes in the graph grows, this can become excessive, especially as we will need more *k*-mers for describing the haplotypes. The solution is compressing the matrices, possibly using methods developed for compressing the color matrices in colored de Bruijn graphs [21]. With a suitable compression scheme, we can also take advantage of the compressed representation to rescore similar haplotypes faster.

We currently sample the haplotypes independently in each block and combine them arbitrarily. This is acceptable with short reads, as the blocks are much longer than the reads. In other applications such as long read mapping, we need phased haplotypes to ensure contiguity across block boundaries. While PanGenie can infer phased haplotypes, the algorithm does not scale well with the number of haplotypes. As a possible solution, we could sample equivalence classes of (almost) identical haplotypes in each block and then choose specific haplotypes from the equivalence classes to maximize contiguity across block boundaries.

With properly phased haplotypes, we may also be able to drop some restrictions on the graph structure. As long as we sample the haplotypes independently in each block, any recombination of them should be a plausible haplotype, because the haplotypes are all constrained to visit the same blocks in order and match each other at the boundaries. But if we can infer phased haplotypes, we can use the long-distance information in them to ensure plausibility instead of relying on these constraints.

Structural variants have been notoriously difficult to study due to their complex and repetitive nature [9], especially so when using the linear reference in combination with short reads [27, 22, 4, 8, 37]. Substantial improvements have been made with the advent of long-read sequencing [17, 15, 32, 20]. However, long reads are comparatively expensive and often unavailable for large cohorts of human samples, such as those underpinning global-scale sequencing projects. The approach we presented, which uses knowledge from population sequencing (and therefore long reads), improves upon the state of the art for short-read SVs genotyping, allowing the typing of common SV variants with accuracy comparable to long-read methods.

Looking forward, we expect pangenomes from the HPRC and elsewhere to substantially grow in genome number, perhaps into the thousands of haplotypes over the next several years. Methods for personalizing pangenomes will therefore increase in importance as the fraction of all variation that is rare within them naturally expands. Conversely, as pangenomes grow the expanding fraction of rare variation they cover should allow increasingly accurate personalized pangenomes to be imputed, providing a framework for genome imputation to include complex structural variation. As a consequence, we expect the relevance and power of the personalization methods of the type introduced here to become increasingly vital to pangenome workflows.

## Acknowledgements

This work was supported in part by the National Human Genome Research Institute (NHGRI) and the National Institutes of Health. B.P. was partly supported by NIH grants: R01HG010485, U24HG010262, U24HG011853, OT3HL142481, U01HG010961, and OT2OD033761.

