## Supplemental material for "Personalized Pangenome References"

December 15, 2023

#### 1 Data sources

##### 1.1 Graphs

We used year-1 HPRC graphs [6] built from 90 haplotypes with the Minigraph–Cactus pipeline [4]. The graphs can be downloaded from `https://github.com/human-pangenomics/hpp_pangenome_resources`. Direct links for the specific graphs (version 1.0 or 1.1, default or frequency-filtered, based on GRCh38 or CHM13) are the following:

- v1.0 default (GRCh38): `https://s3-us-west-2.amazonaws.com/human-pangenomics/pangenomes/freeze/freeze1/minigraph-cactus/hprc-v1.0-mc-grch38.gfa.gz`
- v1.0 frequency-filtered (GRCh38): `https://s3-us-west-2.amazonaws.com/human-pangenomics/pangenomes/freeze/freeze1/minigraph-cactus/filtered/hprc-v1.0-mc-grch38-minaf.0.1.gfa.gz`
- v1.0 default (CHM13): `https://s3-us-west-2.amazonaws.com/human-pangenomics/pangenomes/freeze/freeze1/minigraph-cactus/hprc-v1.0-mc-chm13.gfa.gz`
- v1.0 frequency-filtered (CHM13): `https://s3-us-west-2.amazonaws.com/human-pangenomics/pangenomes/freeze/freeze1/minigraph-cactus/filtered/hprc-v1.0-mc-chm13-minaf.0.1.gfa.gz`
- v1.1 default (GRCh38): `https://s3-us-west-2.amazonaws.com/human-pangenomics/pangenomes/freeze/freeze1/minigraph-cactus/hprc-v1.1-mc-grch38/hprc-v1.1-mc-grch38.gbz`
- v1.1 frequency-filtered (GRCh38): `https://s3-us-west-2.amazonaws.com/human-pangenomics/pangenomes/freeze/freeze1/minigraph-cactus/hprc-v1.1-mc-grch38/hprc-v1.1-mc-grch38.d9.gbz`
- v1.1 default (CHM13): `https://s3-us-west-2.amazonaws.com/human-pangenomics/pangenomes/freeze/freeze1/minigraph-cactus/hprc-v1.1-mc-chm13/hprc-v1.1-mc-chm13.gbz`
- v1.1 frequency-filtered (CHM13): `https://s3-us-west-2.amazonaws.com/human-pangenomics/pangenomes/freeze/freeze1/minigraph-cactus/hprc-v1.1-mc-chm13/hprc-v1.1-mc-chm13.d9.gbz`

##### 1.2 Reads

We used Illumina Novaseq, and Element sequencing data from the following links:

- Novaseq wgs pcr-free data: `gs://brain-genomics-public/research/sequencing/fastq/novaseq/wgs_pcr_free`
- Element wgs: `gs://brain-genomics-public/research/element/cloudbreak_wgs`

##### 1.3 Other data

For benchmarking small variants, we used *Genome in a Bottle* (GIAB) v4.2.1 GRCh38 benchmark sets from the following links:

- HG001 benchmark: `https://ftp-trace.ncbi.nlm.nih.gov/ReferenceSamples/giab/release/NA12878_HG001/NISTv4.2.1/GRCh38/`
- HG002 benchmark: `https://ftp-trace.ncbi.nlm.nih.gov/ReferenceSamples/giab/release/AshkenazimTrio/HG002_NA24385_son/NISTv4.2.1/`
- HG003 benchmark: `https://ftp-trace.ncbi.nlm.nih.gov/ReferenceSamples/giab/release/AshkenazimTrio/HG003_NA24149_father/NISTv4.2.1/GRCh38/`
- HG004 benchmark: `https://ftp-trace.ncbi.nlm.nih.gov/ReferenceSamples/giab/release/AshkenazimTrio/HG004_NA24143_mother/NISTv4.2.1/GRCh38/`
- HG005 benchmark: `https://ftp-trace.ncbi.nlm.nih.gov/ReferenceSamples/giab/release/ChineseTrio/HG005_NA24631_son/NISTv4.2.1/GRCh38/`

For benchmarking structural variants, we used *Genome in a Bottle* (GIAB) Tier1 v0.6 confident regions lifted to GRCh38 annotation as in [6], Challenging Medically Relevent Genes (CMRG) v1.0 and a whole genome BED file from [5] where centromeric regions were removed.

- HG002 benchmark Tier1 v0.6: `https://ftp-trace.ncbi.nlm.nih.gov/ReferenceSamples/giab/release/AshkenazimTrio/HG002_NA24385_son/NIST_SV_v0.6/`
- HG002 benchmark CMRG v1.0: `https://ftp-trace.ncbi.nlm.nih.gov/ReferenceSamples/giab/release/AshkenazimTrio/HG002_NA24385_son/CMRG_v1.00/GRCh38/StructuralVariant/`

#### 2 Selecting sampling parameters

We searched for good haplotype sampling parameters by iterating an early version of the variant calling pipeline with 30x NovaSeq reads for HG002. The graph we used was an unreleased Minigraph–Cactus HPRC graph with some but not all of the improvements in the v1.1 graph. We evaluated the results by comparing the F1 scores separately for indels and SNPs, with the frequency-filtered graph as the baseline. See Supplementary Table 1 for the progression of the F1 scores. Unless otherwise mentioned, all graphs with sampled haplotypes also include reference paths.

First we tried various combinations of $k$-mer scoring parameters while sampling 8 haplotypes. After three rounds and 68 combinations (see Supplementary Table 2), we selected 0.9 as the multiplicative discount for homozygous $k$-mers, 0.05 as the additive adjustment for heterozygous $k$-mers,

| Graph | Stage | Haplotypes | Indel F1 | SNP F1 |
|---|---|---|---|---|
| Frequency-filtered | Baseline | – | 0.993805 | 0.997373 |
| Sampled | $k$-mer scoring (indels) | 8 | 0.993961 | 0.997667 |
| Sampled | $k$-mer scoring (SNPs) | 8 | 0.993935 | 0.997679 |
| Sampled | Number of haplotypes | 4 | 0.994057 | 0.997779 |
| Sampled | Number of haplotypes | 6 | 0.994048 | 0.997786 |
| Sampled | Diploid sampling | 16 | 0.994327 | 0.997891 |
| Sampled | Diploid sampling | 32 | 0.994336 | 0.997891 |

Supplementary Table 1: Progression of F1 scores in various stages of parameter search. For diploid sampling, the number of haplotypes indicates the number of candidates.

| Homozygous | Heterozygous | Absent | Winner | Indel | SNP |
|---|---|---|---|---|---|
| { 0.6, 0.7, 0.8 } | { 0.1, 0.2, 0.3 } | { 0.6, 0.7, 0.8 } | $(0.8, 0.1, 0.8)$ | 0.993817 | 0.997532 |
| { 0.8, 0.85, 0.9 } | { 0, 0.05, 0.1 } | { 0.8, 0.85, 0.9 } | $(0.9, 0.05, 0.8)$ | 0.993961 | 0.997667 |
| { 0.85, 0.9, 0.95 } | { 0, 0.05, 0.1 } | { 0.75, 0.8, 0.85 } | $(0.9, 0.05, 0.8)$ | 0.993961 | 0.997667 |

Supplementary Table 2: Combinations of $k$-mer scoring parameters. For each round, we list the discounts for homozygous $k$-mers, adjustments for heterozygous $k$-mers, scores for absent $k$-mers, the winning combination, and the F1 scores for indels and SNPs with the winning parameters.

and 0.8 as the score for absent $k$-mers. The winning combination had the highest F1 score with indels, while $(0.95, 0.05, 0.85)$ would have been marginally better with SNPs.

Next we locked the $k$-mer scoring parameters and tried sampling $n \in \{2, 4, 6, 8, 10, 12, 14, 16\}$ haplotypes. We found that $n = 4$ was the best choice for indels and $n = 6$ for SNPs. Again, the differences between the F1 scores of the winning values were marginal.

Then we validated the parameter choices by sampling $n \in \{4, 6, 8\}$ haplotypes for HG003, HG004, and HG005 and compared the results to the frequency-filtered graph. We used 30x NovaSeq reads for HG003 and HG004 and 20x, 30x, and 50x reads for HG005. In all cases, sampling either 4 or 6 haplotypes was the best choice.

Finally we tried diploid sampling from $n \in \{4, 8, 16, 32\}$ candidates for HG002. Sampling from 32 candidates was the best choice for indels, while $n = 16$ and $n = 32$ were equally good for SNPs. We also tried sampling from 32 candidates without including reference paths, but the variant calling performance was worse than with reference paths.

### 3 Graphs Properties

| Graph | Reference | Nodes | Edges | Length | Tips | Top-Level Chains |
|---|---|---|---|---|---|---|
| v1.0 | GRCh38 | 81751614 | 113258931 | 3287932785 | 10568 | 2534 |
| v1.0 filtered | GRCh38 | 60243196 | 72669397 | 3153421135 | 46019 | 2698 |
| v1.1 | GRCh38 | 80382908 | 111251520 | 3234968488 | 358 | 173 |
| v1.1 filtered | GRCh38 | 60118570 | 70105101 | 3132180138 | 358 | 173 |

Supplementary Table 3: Properties of the different graphs. Note that there are 12 single-node connected components in the v1.1 graphs, corresponding to small unplaced contigs, which are not counted as "top-level" chains in the vg software.

#### 4 Calling small variants

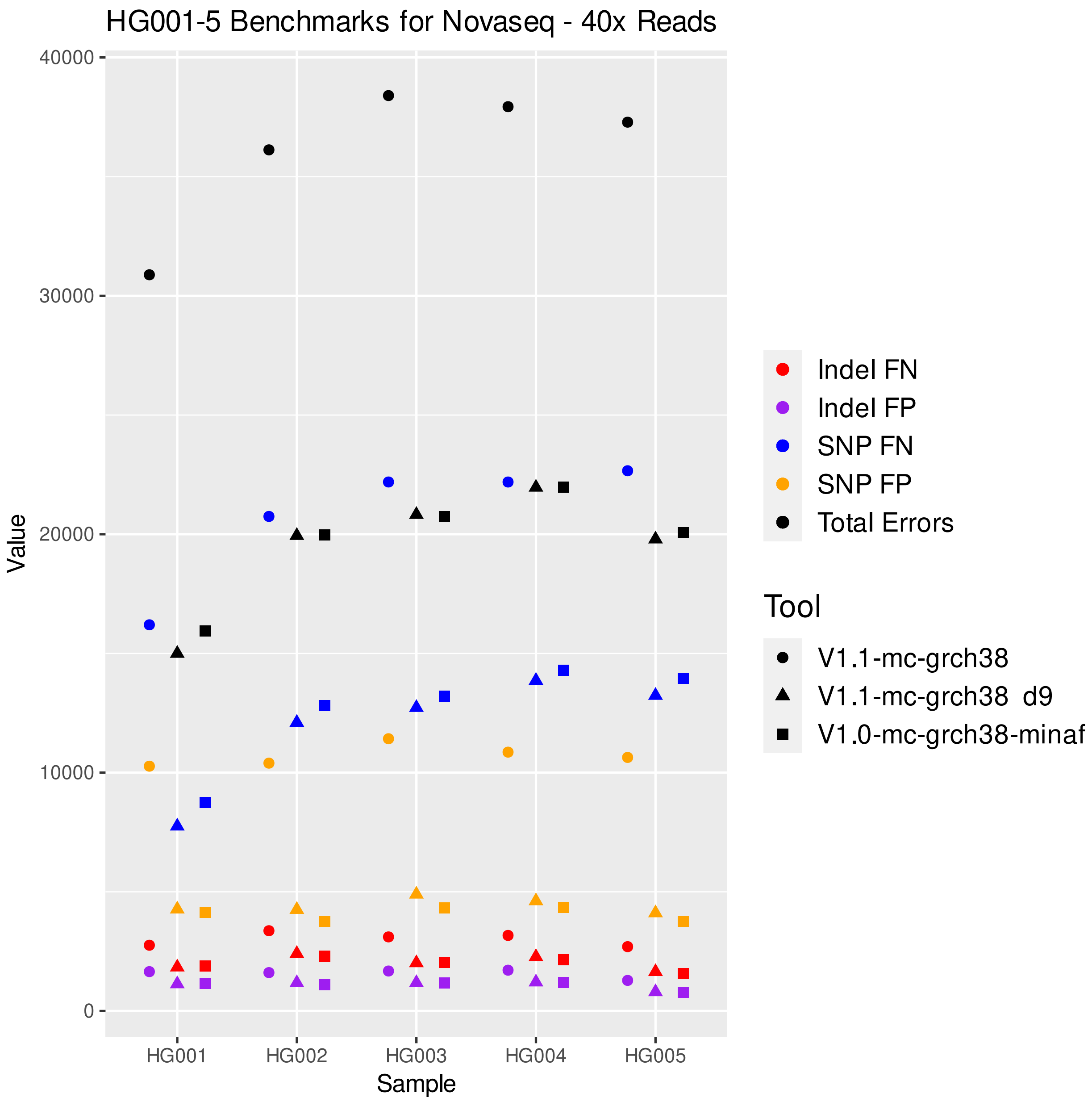


Supplementary Figure 1: Comparing the performance of the frequency-filtered v1.0 and v1.1 graphs and the v1.1 full graph

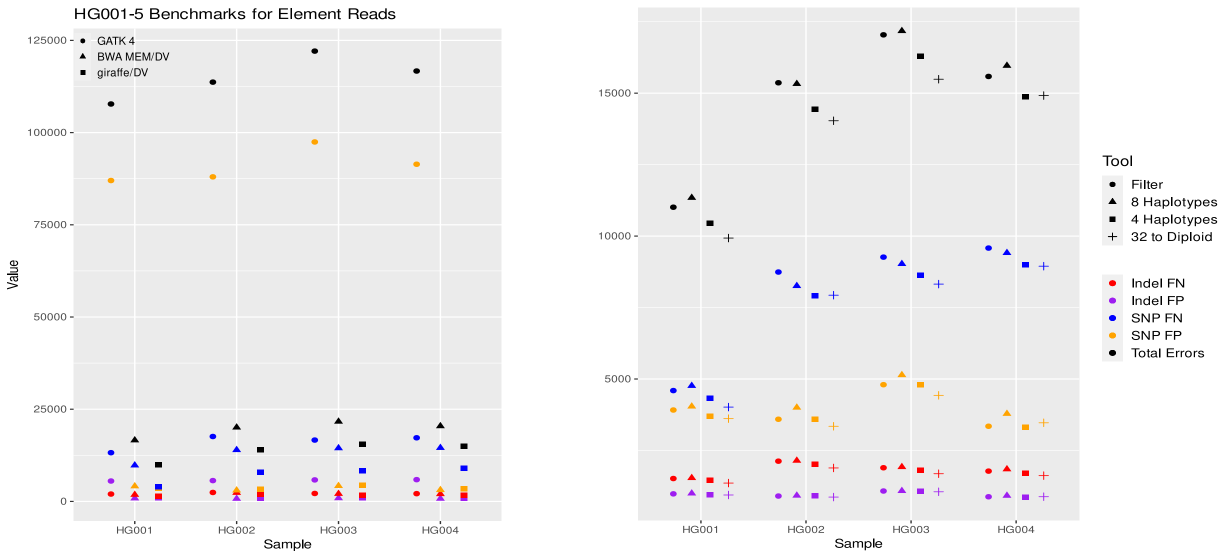


Supplementary Figure 2: Results using Element data. The left panel is comparing the number of errors for different tools, giraffe/DV, BWA MEM/DV, and GATK. The right panel is comparing the results of different graphs used in the giraffe/DV pipeline.

#### 4.1 Command Line

The small variant analysis consist of two main parts before evaluation step.

##### 4.1.1 Haplotype sampling and vg giraffe

We used `vg giraffe` WDL `https://github.com/vgteam/vg_wdl/blob/master/workflows/giraffe.wdl`, and in order to create and use diploid graph in this workflow we used the following flags in running the workflow.

```
HAPLOTYPE_SAMPLING = true
IN_DIPLOID = true
```

We used this workflow to create the Diploid graph (best resulting graph) for the reads, and to map the reads to the Diploid graph. We are using vg 1.49.0 for doing these analysis.

##### 4.1.2 DeepVariant

After having the mapped reads as a .bam file we used the following command to run DeepVariant:

```
docker run\
  google/deepvariant:1.5.0 \
  /opt/deepvariant/bin/run_deepvariant \
  --model_type=WGS \
  --ref=<path_to_reference_file> \
  --reads=<path_to_bam_file> \
  --output_vcf=<path-to-output>.vcf.gz \
  --output_gvcf=<path-to-output>.g.vcf.gz \
  --num_shards=<n_of_threads> \
  --make_examples_extra_args= \
"min_mapping_quality=1,keep_legacy_allele_counter_behavior=true,normalize_reads=true"
```

As seen in the command, we used deepvariant:1.5.0 for our analysis. This step generates .vcf.gz file that is ready for evaluation.

###### 4.1.3 Evaluation using Hap.py

We used hap.py:v0.3.12 for our evaluation analysis using the following command:

```
docker run \
  jmcdani20/hap.py:v0.3.12 /opt/hap.py/bin/hap.py \
  <path_to_truth_set.vcf.gz> \
  <path_to_deepvariant_output>.vcf.gz \
  -f <path_to_truth_set.bed> \
  -r <path_to_reference_file> \
  -o <path_to_happy_output_directory>/happy.output \
  --engine=vcfeval \
  --pass-only
```

#### 5 Genotyping structural variants

For genotyping structural variants (SVs), we used two tools `vg call` v1.49.0 [3] and PanGenie v2.5.0 [1] not only to compare the two but also to validate the robustness of the various rounds of evaluation for the different graphs and call sets we tested for. The main distinction between the two lies in the model adopted to detect SVs. In fact, `vg call` leverages information from reads alignment performed with Giraffe [7] for then augmenting the output file and, finally, calling SVs according to the reads' coverage derived from Giraffe; a more in-depth explanation of the entire process is available in the respective papers [7, 3]. PanGenie adopts an HMM to infer the best possible combination of haplotype paths within the pangenome graph imputed from the short reads' dataset for the sample in question. The resulting genome inference, in VCF format, has to be converted into biallelic sites for the subsequent evaluation to work best; more details on how to run PanGenie can be found on the GitHub page for the tool (PanGenie git) as well as in the corresponding paper [1].

In both cases, the output VCF for the HG002 sample was later evaluated with truvari v3.5.0 [2] exact parametrization, as in [6], optimized for benchmarking using pangenome graphs. This was done consistently across all series of experiments with pangenomes and personalized graphs, truth sets from the GIAB and the T2T consortium and with or without BED regions.

##### 5.1 Command line

These are the commands to generate viable VCF files with `vg call` and PanGenie, respectively. Next, truvari was used for SVs evaluation.

###### 5.1.1 vg call

After reads mapping with `vg giraffe`, the `vg call` workflow includes two additional steps:

```
# outputs an agumented pack file
vg pack -x <graph>.gbz -g HG002.gam -Q 5 -t <n_of_threads> -o HG002.pack
# calls variants for the sample
vg call <graph>.gbz -k HG002.pack -S GRCh38 -t <n_of_threads> -z >
↪  HG002_vgcall.vcf
```

##### 5.1.2 PanGenie

Conversely, PanGenie requires filtering and decomposition of the graph VCF as shown below:

```
# generates the graph VCF if not done already
vg deconstruct -P GRCh38 -a <graph>.gbz -t <n_of_threads> > HG002_graph.vcf
# normalizes the VCF and decomposes large SNARLs
vcfbub -l 0 -a 100000 -i pangenie_ready.vcf.gz | vcfwave -I 10000 -t <n_of_threads> -n > decomposed_and_filtered.vcf
```

However, to ensure the former steps can be executed properly it is necessary to remove haploid sequences form the graph VCF (e.g. CHM13) and artificially make each sample a diploid, phased field in the VCF file; this second operation is not needed when using the v1.1 diploid graph.

```
# removes CHM13 from the graph VCF
cut --complement -d$'\t' -f10 HG002_graph.vcf > HG002_no_chm13.vcf
# splits recombinant haplotypes
awk -F '\t' '/^#/ {print;next;} {OFS="\t"; gsub("[|]","\t",$10)}1' HG002_no_chm13.vcf > HG002_pangenie-spl.vcf
# duplicates recombinant haplotypes and adds them at the end of the VCF file (example for the 4 haplotypes graph)
awk -F '\t' '/^#/ {print;next;} {OFS="\t"; print $0, $10 "\t" $11 "\t" $12 "\t" $13}' HG002_pangenie-spl.vcf > HG002_pangenie-dup.vcf
# reorganizes columns for each diploid pair to be a perfect copy of each other (example for the 4 haplotypes graph)
awk -F '\t' '/^#/ {print;next;} {OFS="\t"; print $1, $2, $3, $4, $5, $6, $7, $8, $9, $10, $14, $11, $15, $12, $16, $13, $17}' HG002_pangenie-dup.vcf > HG002_input-phased.vcf
# phases the VCF
awk -F '\t' '/^#/ {print;next;} {OFS="\t";for(i=j=10; i < NF; i+=2) {$j = $i"|"$(i+1); j++} NF=j-1}1' HG002_input-phased.vcf > pangenie_ready.vcf
# renames each recombinant sample for PanGenie to interpret the actual number of samples in the VCF
vim pangenie_ready.vcf #manually edit the column headers to match the number of columns in file
```

Afterwards, PanGenie can be run in pipeline mode, passing through Snakemake, so that the resulting VCF file has a structure compatible with the script for biallelic conversion. This entails having a working CONDA version (Anaconda Software Distribution. Computer software. Vers. 3-23.3.1. Anaconda, Mar. 2023. Web.) with Mamba installed (version used 1.2.0) which is mandatory to run Snakemake (version used 7.30.1). Both PanGenie and Snakemake have to be placed in a separate CONDA environment and upon installing and building the tool, at this location *path/to/pangenie/pipelines/run-from-callset*, it will be present a config.yaml file to be filled in with relevant information. Once done, PanGenie can be run after activating the Snakemake environment and invoking the following from the folder where the config.yaml is saved:

```
# runs PanGenie
snakemake --cores <n_of_threads>
```

In general, the config.yaml file will ask for a graph VCF generated as previously described, a reference file (GRCh38 in this case), the reads dataset for the sample in question (HG002 in this set of analyses), the paths to the binaries for the tool and an output directory. Chromosomes' name must be identical both in the reference and in the graph VCF for the genome inference to be performed correctly; it is good practice to leave the reference untouched and change the name of chromosomes in the VCF file with this single command line:

```
# renames chromosomes in VCF (based on a GRCh38 graph)
sed -i 's/GRCh38#0#//' decomposed_and_filtered.vcf
```

Further details on how to run PanGenie can be found on the GitHub page of the tool, where there are also specifications on how to carry out the biallelic conversion with an ad-hoc Python script. To be noted, PanGenie output will be stored in a folder at the location *path/to/output/directory/genotypes*; this is the VCF file benefiting from biallelic conversion before evaluation.

##### 5.1.3 Variants evaluation

At last, variants evaluation is done running truvari like so:

```
# SVs evaluation
truvari bench -b <truth_set>.vcf.gz -c <compare>.vcf.gz -f
↪ <path_to_reference_file> --includebed <path_to_bed_file> -O 0.0 -r 1000 -p
↪ 0.0 -P 0.3 -C 1000 -s 50 -S 15 --sizemax 100000 --multimatch --no-ref c -o
↪ <output_directory>
```

where the option for including a BED file can be toggled on or off whether or not the analysis conducted was using it; other parameters were left the same as [6] to attain the best performance when benchmarking against pangenome graphs.

| **Strategy** | **Task** | **Runtime** |
|---|---|---|
| PanGenie-GRCh38 | filtering and decomposition | 5117 |
| PanGenie-GRCh38 | genome inference | 5112 |
| `vg call`-GRCh38 | pack file | 4365 |
| `vg call`-GRCh38 | calling variants | 2741 |

Supplementary Table 4: Runtime comparisons for both PanGenie and `vg call` on the v1.1 diploid graph.

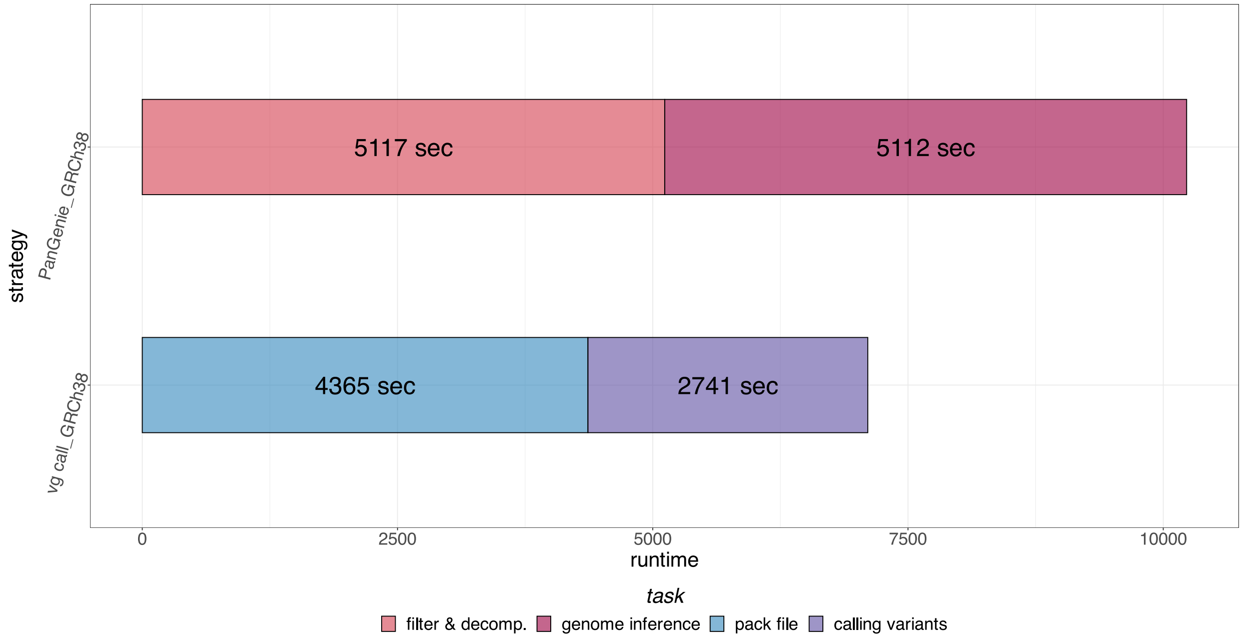


Supplementary Figure 3: Runtimes divided by tasks for the PanGenie and `vg call` v1.1 diploid graph structural variants evaluation. Filtering and decomposition is required to effectively run the PanGenie genome inference task, whereas a pack file needs to be generated before preforming variant calling with `vg call`. Reads alignment time by `vg giraffe` functional to variant calling by `vg call` not shown.
